# Cocaine-induced locomotor activation differs across six sets of inbred mouse substrains

**DOI:** 10.1101/2021.11.08.467748

**Authors:** Christiann H. Gaines, Sarah A. Schoenrock, Joseph Farrington, David F. Lee, Lucas J. Aponte-Collazo, Ginger Shaw, Darla R. Miller, Martin T. Ferris, Fernando Pardo-Manuel de Villena, Lisa M. Tarantino

**Affiliations:** Department of Genetics, School of Medicine, University of North Carolina at Chapel Hill, NC, United States; Neuroscience Curriculum, University of North Carolina at Chapel Hill, NC, United States; Pharmacology Curriculum, University of North Carolina at Chapel Hill, NC, United States; Department of Pharmacology, School of Medicine, University of North Carolina at Chapel Hill, NC, United States; Lineberger Comprehensive Cancer Center, School of Medicine, University of North Carolina at Chapel Hill, NC, United States; Division of Pharmacotherapy and Experimental Therapeutics, Eshelman School of Pharmacy, University of North Carolina at Chapel Hill, NC, United States

**Keywords:** cocaine sensitivity, initial cocaine response, genetics, reduced complexity cross, rodent behavior, addiction, rodent model, mice models

## Abstract

Cocaine use disorders (CUD) are devastating for affected individuals and impose a significant burden on society, but there are currently no FDA-approved therapies. The development of novel and effective treatments has been hindered by substantial gaps in our knowledge about the etiology of these disorders. The risk for developing a CUD is influenced by genetics, the environment and complex interactions between the two. Identifying specific genes and environmental risk factors that increase CUD risk would provide an avenue for the development of novel treatments.

Rodent models of addiction-relevant behaviors have been a valuable tool for studying the genetics of response to drugs of abuse. Traditional genetic mapping using genetically and phenotypically divergent inbred mice has been successful in identifying numerous chromosomal regions that influence addiction-relevant behaviors, but these strategies rarely result in identification of the causal gene or genetic variant. To overcome this challenge, reduced complexity crosses (RCC) between closely related inbred mouse substrains have been proposed as a method for rapidly identifying and validating functional variants. The RCC approach is dependent on identifying phenotypic differences between substrains. To date, however, the study of addiction-relevant behaviors has been limited to very few sets of substrains, mostly comprising the C57BL/6 lineage.

The present study expands upon the current literature to assess cocaine-induced locomotor activation in 20 inbred mouse substrains representing six inbred strain lineages (A/J, BALB/c, FVB/N, C3H/He, DBA/2 and NOD) that were either bred in-house or supplied directly by a commercial vendor. To our knowledge, we are the first to identify significant differences in cocaine-induced locomotor response in several of these inbred substrains. The identification of substrain differences allows for the initiation of RCC populations to more rapidly identify specific genetic variants associated with acute cocaine response. The observation of behavioral profiles that differ between mice generated in-house and those that are vendor-supplied also presents an opportunity to investigate the influence of environmental factors on cocaine-induced locomotor activity.

## 1 Introduction

Cocaine remains the second most commonly used drug worldwide and both cocaine use and cocaine related overdose deaths have been increasing in the United States in recent years (Karila, Petit et al. 2012, Jalal, Buchanich et al. 2018, Bentzley, Han et al. 2021). Over half of cocaine users in the United States have a diagnosed cocaine use disorder (CUD), indicating the high addiction liability of cocaine (Goldman, Oroszi et al. 2005, SAMHSA 2019). Despite the prevalence of CUD, there are currently no FDA approved therapies. The lack of treatment options is partially due to gaps in our knowledge about the etiology of this complex and devastating disorder.

Not all who use cocaine will go on to develop a CUD, suggesting that there are individual differences in risk. The risk of developing a CUD is influenced by genetics, the environment and interactions between the two (Merikangas and Avenevoli 2000, Goldman, Oroszi et al. 2005, Thatcher and Clark 2008, Ducci and Goldman 2012, Mennis, Stahler et al. 2016). Twin studies suggest heritability estimates of approximately 0.70 for cocaine dependence indicating a significant contribution of genetics (Ducci and Goldman 2012). Human genome wide association studies (GWAS) are currently being conducted to identify specific loci and genes associated with substance use disorders (Sullivan, Agrawal et al. 2018) and have been highly successful for some drugs of abuse such as nicotine and alcohol. However, very few GWAS studies have been published for cocaine dependence or CUD (Gelernter, Sherva et al. 2014, Cabana-Dominguez, Shivalikanjli et al. 2019, Sun, Kranzler et al. 2020) due to lack of sufficient sample size of individuals with cocaine dependence for current GWAS methods, which has limited the genetic discovery in GWAS for specific loci and genes. Identifying genes and molecular pathways that increase CUD risk would provide insight into individuals at increased risk and novel targets that could be investigated for the development of therapeutics.

A complementary approach to human GWAS studies of cocaine use and development of CUD is genetic mapping studies and mechanistic follow-up of GWAS loci in rodent models. A notable example is the replication of the identification of family with sequence similarity 53, member B (*FAM53B*), as a risk variant for cocaine dependence in a mouse mapping population for self-administration of cocaine (Gelernter, Sherva et al. 2014, Dickson, Miller et al. 2016). The use of rodent models offers several advantages, including the ability for the genetic background, environment, and drug exposure regimens to be controlled and precisely manipulated. While rodent models cannot fully recapitulate the range of symptoms observed in humans with CUD, they do allow for measurement of specific addiction-relevant behaviors, including initial drug sensitivity. Retrospective and longitudinal studies in humans have shown that individual differences in initial subjective drug responses can predict subsequent drug use (Haertzen, Kocher et al. 1983, Davidson, Finch et al. 1993, Lambert, McLeod et al. 2006, de Wit and Phillips 2012). In mice, acute locomotor response to an initial dose of cocaine is a well-established model of initial sensitivity (Thomsen and Caine 2011, Wiltshire, Ervin et al. 2015).

Traditional genetic mapping approaches using inbred mouse strains have been used successfully to identify genomic regions, termed quantitative trait loci (QTL), that are associated with cocaine-induced locomotor activation (Tolliver, Belknap et al. 1994, Miner and Marley 1995, Phillips, Huson et al. 1998, Jones, Tarantino et al. 1999, Boyle and Gill 2001, Gill and Boyle 2003, Boyle and Gill 2009, Dickson, Miller et al. 2016). Mapping approaches involve outcrosses between genetically and phenotypically diverse pairs of inbred strains and the resulting F1s are intercrossed or backcrossed to generate F2 or N2 mapping populations, respectively. The resulting QTL identified from a F2 or N2 population typically span tens of megabases and comprise hundreds of genes and thousands of potential causal polymorphisms. Therefore, identifying the causative variants that affect cocaine-induced locomotor activation and other complex behavioral traits has been extremely challenging.

The use of Reduced Complexity Crosses (RCC) for genetic mapping offers a significant advantage over traditional genetic mapping strategies. Mouse substrains are nearly isogenic inbred mouse substrains that are derived from the same founder inbred strain but have been bred independently for many generations (typically more than 20). An RCC is generated in the same fashion as an F2 or N2 population described above, using two substrains that differ for a phenotype of interest. The size of the resulting QTL confidence interval is similar to a traditional F2 mapping population, spanning ten(s) of megabases. However, within a QTL interval, the two nearly isogenic substrains will have significantly fewer variants, limited to polymorphisms that were either segregating at the time the strains were separated or arose spontaneously after that time (Bryant CD 2018). This feature dramatically facilitates follow-up on the few polymorphisms within the QTL and identification of the causative polymorphism. Additionally, a recently developed genotyping array captures polymorphisms between inbred mouse substrains, facilitating rapid and reliable genotyping of RCCs (Sigmon, Blanchard et al. 2020). A RCC has been used successfully to identify a genetic polymorphism in the *Cyfip2* gene that results in differing psychostimulant responses in C57BL/6J and C57BL/6N substrains (Kumar, Kim et al. 2013).

Genetic differences between substrains are likely to influence any number of phenotypes, offering a powerful tool with which to expand our knowledge about the genetic loci that affect addiction-relevant behaviors. Thus far, the literature describing substrain differences in locomotor response to drugs of abuse has focused on C57BL/6 substrains. In this study, we measured cocaine-induced locomotor activation across 6 substrains derived from A/J, BALB/c, DBA/2, FVB/N, NOD and C3H/He inbred mouse strains. We report significant differences across these strains within their respective substrains for acute cocaine sensitivity. These data significantly expand knowledge about substrain differences in cocaine locomotor response and offer the opportunity to pursue genetic studies to identify genes that contribute to this behavior.

## 2 Materials and Methods

### 2.1 General Methods

Mice were all housed in a pathogen-free facility at UNC. This facility consisted of a 12-hour light/dark cycle with lights on at 7:00 AM. All animal care and protocols were approved by the The University of North Carolina at Chapel Hill (UNC) Institutional Animal Care and Use Committees and followed guidelines that were implemented by the National Institutes of Health Guide for the Care and Use of Laboratory Animals., 8^th^ Edition. Mice were maintained in AAALAC-accredited, specific pathogen free (SPF) barrier colony in ventilated cages (Tecniplast, Buguggiate, Italy). Food (PicoLab Rodent Diet 20, Purina, St. Louis, Missouri) was provided *ad libitum* and throughout the duration of behavioral testing. Edstrom carbon filtered; reverse osmosis hyper-chlorinated water was provided *ad libitum* except during behavioral testing.

Two groups of mice were used for behavioral testing (see **Supp Table 1** for a summary of substrains, origin, housing, and vendor). The first group consisted of five sets of substrains that were purchased directly from their respective commercial vendors and were then group housed by substrain and sex. The second group consisted of five sets of substrains that were originally purchased from their respective vendors, but mice tested in this study were bred in our laboratory at UNC. Mice bred at UNC were either group-housed with cagemates of the same substrain or co-housed with mice from other substrains within their strain group (i.e. DBA/2J, DBA/2NCrl and DBA/2NTac mice in the same cage). Mice were co-housed at weaning, around postnatal day 21.

Vendor supplied substrains were an average age of 62 days old at the start of testing. In-house bred mice were an average age of 65 days old at the start of testing. All mice were weighed on the day prior to testing and weights were used to determine the volume of saline or cocaine administered during open field testing. On the day of testing, mice were transported to the testing room, which was located within the same vivarium, immediately prior to the start of testing. Behavioral testing occurred during the light cycle from 8:00 AM to 12:00 PM with the time that a mouse was tested being consistent across the three test days.

### 2.2. Animals

#### 2.2.1. Vendor Supplied Substrains

A/J, BALB/c, FVB/N and DBA/2 substrains were purchased from their respective commercial vendors and housed in substrain specific cages throughout testing. Mice were an average of 27 days old upon arrival to UNC and were acclimated to the vivarium for 5 weeks after arrival before behavioral testing. A/J substrains were A/J (The Jackson Laboratory, 000646), A/JCr (Charles River Laboratories, 563) and A/JOlaHsd (Envigo, 049). BALB/c substrains were BALB/cJ (The Jackson Laboratory, 000651), BALB/cByJ (The Jackson Laboratory, 001026), BALB/cAnNCrl (Charles River Laboratories, 028), BALB/cAnNHsd (Envigo, 047). FVB/N substrains were FVB/NJ (The Jackson Laboratory, 001800), FVB/NCrl (Charles River Laboratories, 207), FVB/NHsd (Envigo, 118), and FVB/NTac (Taconic Biosciences, FVB-F/FVB-M). DBA/2 substrains were DBA/2J (The Jackson Laboratory, 000671), DBA/2NCrl (Charles River Laboratories, 026) and DBA/2NTac (Taconic Biosciences, DBA2-F/DBA2-M).

#### 2.2.2. Substrains Bred In-House

Another cohort of inbred mouse substrains were purchased from commercial vendors but test animals were bred in-house at UNC. The following substrains were tested in this cohort: DBA/2J, DBA/2NCrl, DBA/2NTac, A/J, A/JOlaHsd, BALB/cByJ, BALB/cJ, FVB/NJ, FVB/NTac, NOD/MrkTac (Taconic Biosciences, NOD-F/NOD-M), NOD/ShiLtJ (The Jackson Laboratory, #001976), C3H/HeJ (The Jackson Laboratory, #000659), C3H/HeNTac (Taconic Biosciences, C3H-F/C3H-M), C3H/HeNHsd (Envigo, 040) and C3H/HeNCrl (Charles River Laboratories, 025). Some of these mice were cohoused with cagemates that included at least two different substrains from a single progenitor strain. Information on all inbred substrains including cage environment (cohoused vs not cohoused) is provided in **Supplemental Table 1**.

### 2.3 Drugs

Cocaine hydrochloride (HCl) was purchased from Sigma-Aldrich (St. Louis, MO, USA; C5776-5G). A solution of cocaine HCl was prepared fresh daily. Cocaine HCl was dissolved in physiological saline at a concentration of 2 mg/ml and administered via intraperitoneal (i.p.) injection at a volume of 0.01 ml/g resulting in a dose of 20 mg/kg of body weight administered to mice for behavioral testing. Saline was purchased from Fisher Scientific (Waltham, MA, USA; 297753).

### 2.4 Open Field Apparatus

The open field (OF) arena (ENV-515-16, Med Associates, St. Albans, VT, USA), measured 17×17×13cm and consisted of four clear Plexiglas walls and a white Plexiglas floor. The walls are surrounded by infrared detection beams on the X, Y and Z axes used to detect horizontal and vertical activity of the animal throughout the duration of the test session. The OF chamber is placed within a sound attenuating box (73.5×59×59 cm) that has two overhead light fixtures containing 28-V lamps. Light levels on the arena floor were 24 lux in the center, 10 lux in the corners and 13 lux along the walls. Eight identical OF arenas were used for testing with a mouse being tested in the same arena each test day.

### 2.5 Acute Cocaine-Induced Locomotor Activity Test

On day 1 (habituation) and 2 (baseline), mice were given an i.p. injection of saline at a volume of 0.01 ml/g body weight and immediately placed into the OF chambers for 30 mins. On day 3 (acute cocaine exposure), mice were given an i.p. injection of 20 mg/kg of cocaine and placed into the OF chamber for 30 mins. At the end of each test session, mice were placed back into their home cages and the OF chambers were cleaned with 0.25% bleach. Locomotor behavior was measured as total distance moved (in centimeters) for the entire 30-min test period each day using the manufacturers data acquisition software (Activity Monitor v5.9.725; Med Associates).

### 2.6 Statistical Analysis

All statistical analyses were performed using SPSS v28 for Mac (IBM Inc). For each set of substrains, we performed an ANOVA that included day of testing, substrain and sex as independent variables and locomotor activity as the dependent variable. Significant main effects (*p* < 0.05) were followed up with post-hoc Tukey’s HSD or independent samples T-tests.

## 3 Results

Experimental data for all inbred mouse substrains including origin of the mice, number of mice tested, cage environment, strain means and standard deviations are provided in **Supplemental Table 1**. We observed significant substrain differences in basal and/or cocaine-induced locomotor activity in 4 of the 6 strain groups we examined. C3H/He and DBA/2 substrain differences were fairly stable across experimental cohorts. We also observed substrain differences (i.e. A/J and FVB/N) that were not replicated across experimental groups (vendor-supplied vs in-house). Sex differences also varied across and within strain groups and experimental cohorts. Results presented individually by substrain are described below.

### 3.1 A/J substrain behavior differs across experimental cohorts

#### Vendor Supplied

Overall, the locomotor activity of the vendor-supplied A/J substrains did not increase significantly after exposure to cocaine (F(_2,125_) = 0.82; *p* = 0.445), although increases in cocaine-induced locomotor activity can be observed in A/JCr and A/JOlaHsd substrains (**Fig 1A**). There was a significant main effect of substrain (F(_2,125_) = 5.5; *p* = 0.005). A/JCr mice were significantly more active than either A/J (*p* = 0.047) and A/JOlaHsd mice (*p* = 0.006). No significant sex (F(_1,125_) = 0.05; *p* = 0.827) or interaction effects were observed.

**Figure 1.**
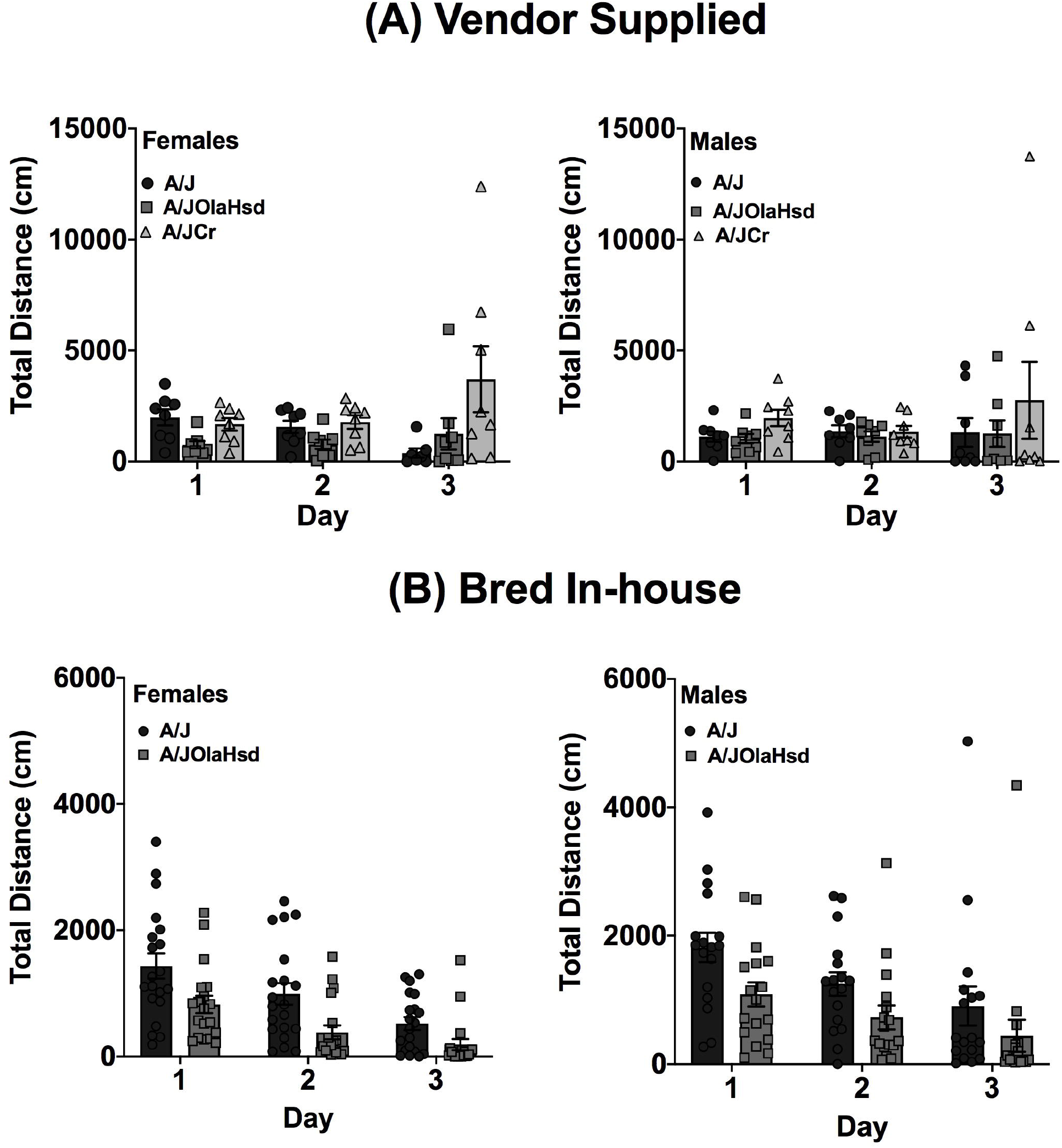
Acute cocaine induced locomotor activity for vendor supplied (not cohoused) and bred in-house (cohoused) A substrains. (A) Total distance moved in 30 mins for each test day for vendor supplied female and male A substrains. While not significant, there was an overall increase in cocaine induced locomotor activity in the A/JCr and A/JOlaHsd substrains. (B) Total distance moved in 30 mins for each test day for bred in-house female and male A substrains. For both A substrains there was a decrease in locomotor activity across test days for male and female mice. Male mice were significantly more active compared to female mice. Each data point represents an individual mouse and error bars are standard error of the mean.

#### Bred In-House

Two A/J substrains were bred in-house at UNC for 1-2 (A/JOlaHsd) or 3-4 (A/J) generations. These mice showed a dramatically different behavioral profile than those obtained from commercial vendors (**Fig 1B**). We observed a significant decrease in locomotor activity across all three days (F_(2,207)_ = 17.8; p = 6.9×10^−8^). We also observed significant substrain (F(_1,207_) = 26.1; *p* = 7.4 × 10^−7^) and sex (F(_1,207_) = 8.6; *p* = 0.004) effects. The A/J substrain showed significantly higher locomotor activity across all three days (*t*(208) = 4.7; *p* = 5.0 × 10^−6^) and males were significantly more active than females (*t*(182) = 2.5; *p* = 0.012). There were no significant interactions among any of the independent variables tested.

### 3.2 Locomotor response to cocaine differs in BALB/c substrains from the “J” lineage in comparison with substrains from the “AnN” lineage

#### Vendor Supplied

We observed significant changes in locomotor activity in the four vendor-supplied BALB/c substrains across three days of testing (F(_2,162_) = 7.7; *p* = 6.2×10^−4^). Collapsed across strains, locomotor activity in response to cocaine on Day 3 is significantly higher than Day 2 (p = 0.006) but not Day 1 (p = 0.889) activity in response to saline. We also observed a significant substrain (F(_3,162_) = 13.7; *p* = 5.0×10^−8^) and substrain x day interaction (F(_6,162_) = 2.7; *p* = 0.017) (**Fig 2**). Mice of both J substrains (BALB/cJ and BALB/cByJ) are significantly more active than BALB/cAnNHsd mice on Day 1 (both *p* < 0.05). Although mice of both J substrains are more active than BALB/cAnNCrl mice on Day 1, the difference is only significant for BALB/cJ mice (*p* < 0.05). The same general pattern is observed for locomotor activity on Day 2 in the J strains vs BALB/cAnNCrl, but neither J substrain differs from BALB/cAnNHsd. Finally, on Day 3 BALB/cByJ mice are significantly more active in response to 20 mg/kg cocaine in comparison to BALB/cAnNCrl and BALB/cAnNHsd (both *p* < 0.01). Female mice were significantly more active than male mice across all days and collapsed across substrain (*t*(184) = −2.0; *p* = 0.023).

**Figure 2.**
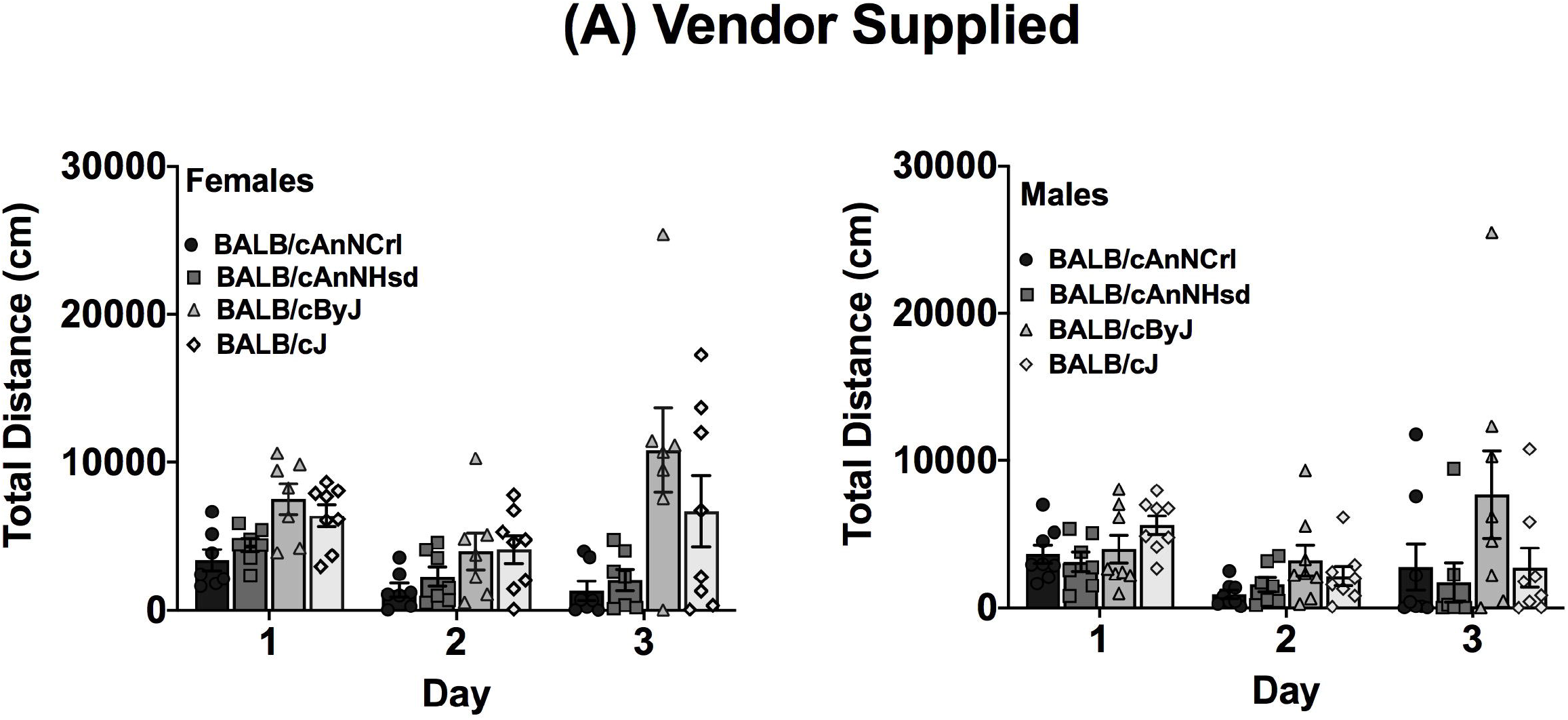
Acute cocaine induced locomotor activity for vendor supplied BALB/c substrains. Total distance moved in 30 mins for each test day for vendor supplied female and male BALB/c substrains. Across all test days female mice had significantly higher locomotor activity compared to male mice. Each data point represents an individual mouse and error bars are standard error of the mean.

#### Bred In-House

BALB/cJ and BALB/cByJ mice were generated in-house and offspring from the 1^st^ and 2^nd^ generations of breeding from the initial vendor stock. We observed no significant day (F(_2,27_) = 0.87; p = 0.431) or substrain (F(_1,27_) = 0.01; p = 0.927) differences (**data not shown**). We also observed no sex differences (F(_1,27_) = 2.3; p = 0.145) but it should be noted that our experimental cohort was limited to only 2 BALB/cByJ males and no BALB/cJ males.

### 3.3 Basal and cocaine-induced locomotor differences are consistent across different experimental cohorts of C3H/He substrains

#### Bred in-house

All C3H/He substrains were bred in-house. The initial cohort was limited to C3H/HeJ and C3H/HeNTac substrains and were the first generation of offspring from vendor-supplied mice (C3H/HeJ) or offspring of crosses between mice from the third generation bred at UNC (C3H/HeNTac). The second cohort of mice included C3H/HeNCrl and C3H/HeNHsd substrains in addition to C3H/HeNTac and C3H/HeJ, and were produced by breeding vendor-supplied mice at UNC for one generation.

Both C3H/HeJ and C3H/HeNTac substrains in the first cohort were significantly more active in response to cocaine (Day 3) vs saline (Days 1 and 2) (F(_2,105_) = 216.0; p = 8.3 × 10^−46^). We also observed a significant substrain effect (F(_1,105_) = 39.3; p = 8.2 × 10^−9^) and substrain x day interaction (F_(2,105)_ = 15.3; *p* = 1.0 × 10^−6^). C3H/HeNTac mice were significantly more active than C3H/HeJ mice in response to saline (**Fig 3A**) on Days 1 (*t*(37) = −5.7; *p* = 2.0 × 10^−6^) and 2 (*t*(37) = −4.2; *p* = 1.6 × 10^−4^) and in response to cocaine on Day 3 (*t*(29) = −5.1; *p* = 1.8 × 10^−5^). No sex differences were observed (F(_1,105_) = 2.3; *p* = 0.132).

**Figure 3.**
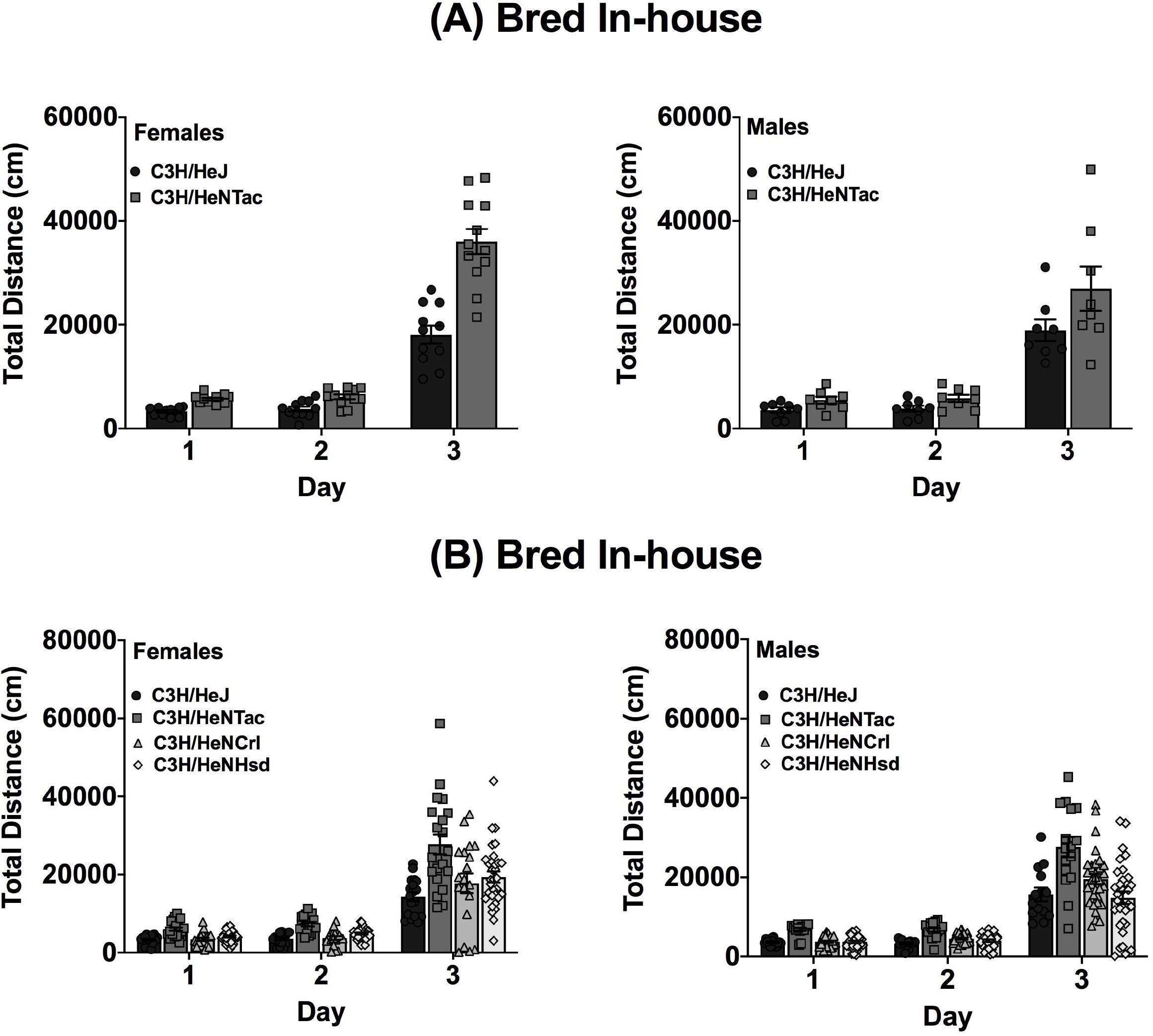
Acute cocaine induced locomotor activity for C3H/He substrains bred in-house. (A) Total distance moved in 30-mins across each test day for cohoused female and male C3H mice. While there were no sex differences, C3H/HeNTac male and female mice had higher locomotor activity and cocaine induced locomotor activity compared to C3H/HeJ mice. Both substrains, regardless of sex, had significantly higher activity following cocaine exposure on day 3 compared to days 1 and 2. (B) Total distance moved in 30-mins across each test day for non-cohoused female and male C3H mice. Male and female C3H/HeNTac mice had higher locomotor activity and cocaine induced locomotor activity compared to the other three substrains. Each data point represents an individual mouse and error bars are standard error of the mean.

In the second cohort comparing all four C3H/He substrains, we observed a significant effect of substrain (F(_3_,_528_) = 32.7; p = 2.2 × 10^−19^) and day (F(_2,528_) = 467.6; *p* = 1.4 × 10^−117^) and a substrain by day interaction (F(_6,528_) = 8.0; *p* = 2.9 × 10^−8^). As in the first cohort, C3H/HeNTac mice were significantly more active than C3H/HeJ (*p* = 5.1 × 10^−13^) and both of the additional C3H/He substrains (**Fig 3B**; all *p* < 0.001). No sex differences were observed (F(_1,528_) = 0.031; *p* = 0.86).

### 3.4 Vendor-supplied and in-house DBA/2 substrains show strikingly similar differences in basal and cocaine-induced locomotor behavior

#### Vendor Supplied

We observed significant day (F(_2,126_) = 23.0; *p* = 3.1×10^−9^) and substrain (F(_2,126_) = 13.4; *p* = 5.0×10^−6^) differences among the three vendor-supplied DBA/2 substrains – DBA/2J, DBA/2NTac and DBA/2NCrl. There was also a significant day x substrain interaction (F(_4,126_) = 3.4; *p* = 0.011). DBA/2NTac mice were significantly less active than DBA/2NCrl mice on Day 1 (*p* = 0.04) and both DBA/2NCrl and DBA/2J mice on Days 2 (all *p* <0.01) and 3 (all *p* < 0.05). No significant sex differences were observed (**Fig 4A**).

**Figure 4.**
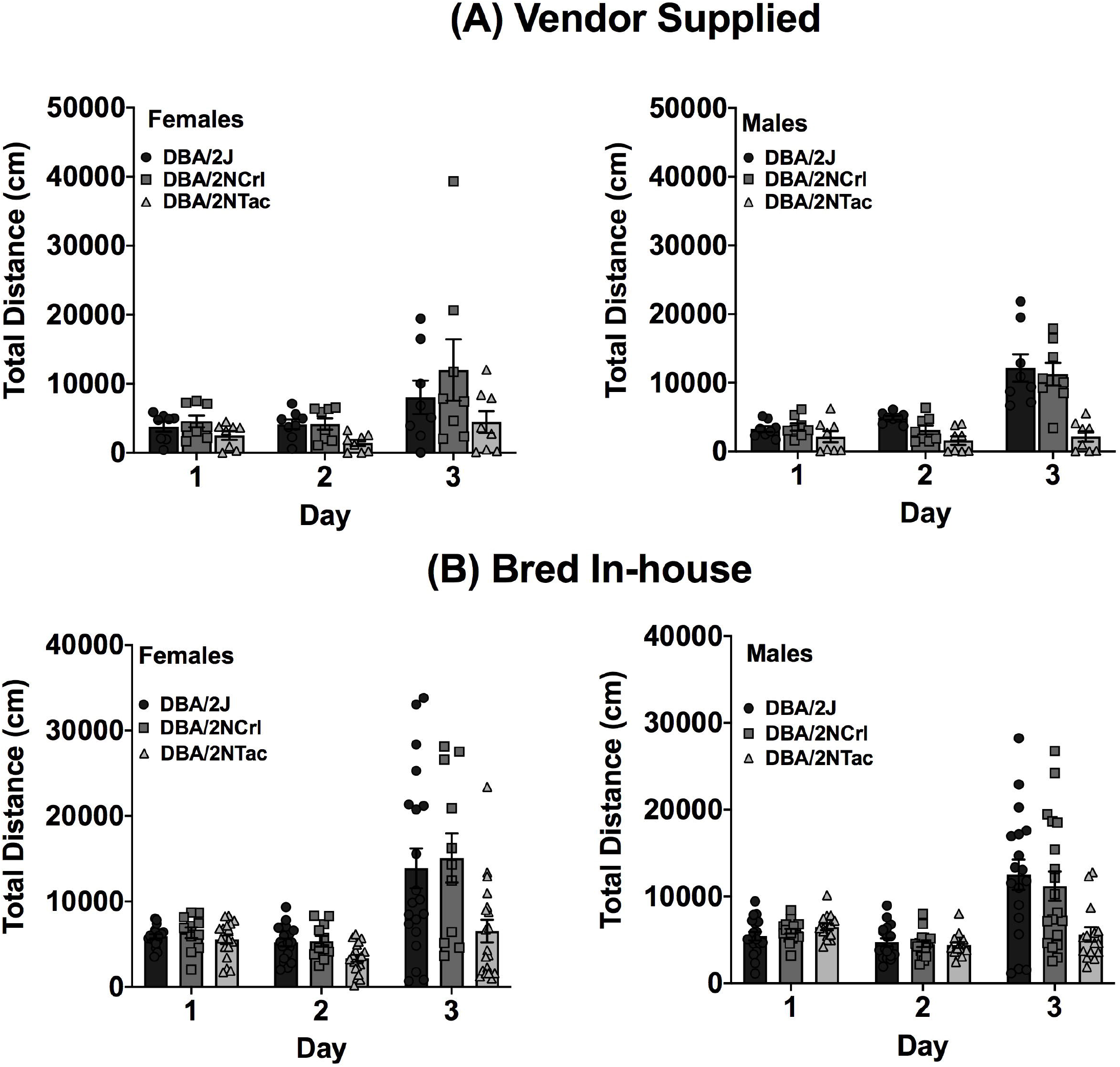
Acute cocaine induced locomotor activity for DBA/2 substrains that were vendor supplied or bred in-house. (A) Total distance moved in 30-mins across each test day for vendor supplied female and male DBA/2 substrains. Both female and male DBA/2 substrains had higher locomotor activity following cocaine exposure compared to activity on days 1 and 2. (B) Total distance moved in 30-mins for each test day for DBA/2 substrains that were bred in-house. Substrains were cohoused at weaning. All female DBA/2 substrains had higher cocaine induced locomotor activity compared to days 1 and 2. All male DBA/2 substrains, except DBA/2NTac, had higher activity following cocaine compared to their activity on days 1 and 2. Each data point represents an individual mouse and error bars are standard error of the mean.

#### Bred In-House

The same set of DBA/2 mice were bred in-house at UNC and mice from the first generation were tested for cocaine-induced locomotor activity. We observed a similar pattern of behavior in these DBA/2 substrains. There were significant day (F(_2,288_) = 48.1; *p* = 9.1×10^−19^) and substrain (F(_2,288_) = 11.0; *p* = 2.4×10^−5^) effects and a significant day x substrain interaction (F(_4,288_) = 7.7; *p* = 6.0×10^−6^). Although none of the substrains differed for locomotor activity on Day 1, DBA/2NTac mice had significantly lower locomotor activity than DBA/2J mice on Day 2 (*p* = 0.018) and both DBA/2J (*p* = 4.6×10^−4^) and DBA/2NCrl mice (*p* = 0.003) on Day 3 (**Fig 4B**).

### 3.5 Basal and cocaine-induced locomotor behavior differs across FVB/N mice bred in-house but not vendor-supplied cohort

#### Vendor Supplied

We observed no significant substrain (F(_3,72_) = 0.76; *p* = 0.519) or sex (F(_1,72_) = 0.03; *p* = 0.859) differences and no significant interaction effects among the four vendor-supplied FVB/N substrains – FVB/NJ, FVB/NTac, FVB/NCrl and FVB/NHsd. However, all FVB/N substrains showed significantly increased locomotor activity on Day 3 after exposure to cocaine (F(_2,72_) = 123.2; *p* = 5.7×10^−24^; **Fig 5A**).

**Figure 5.**
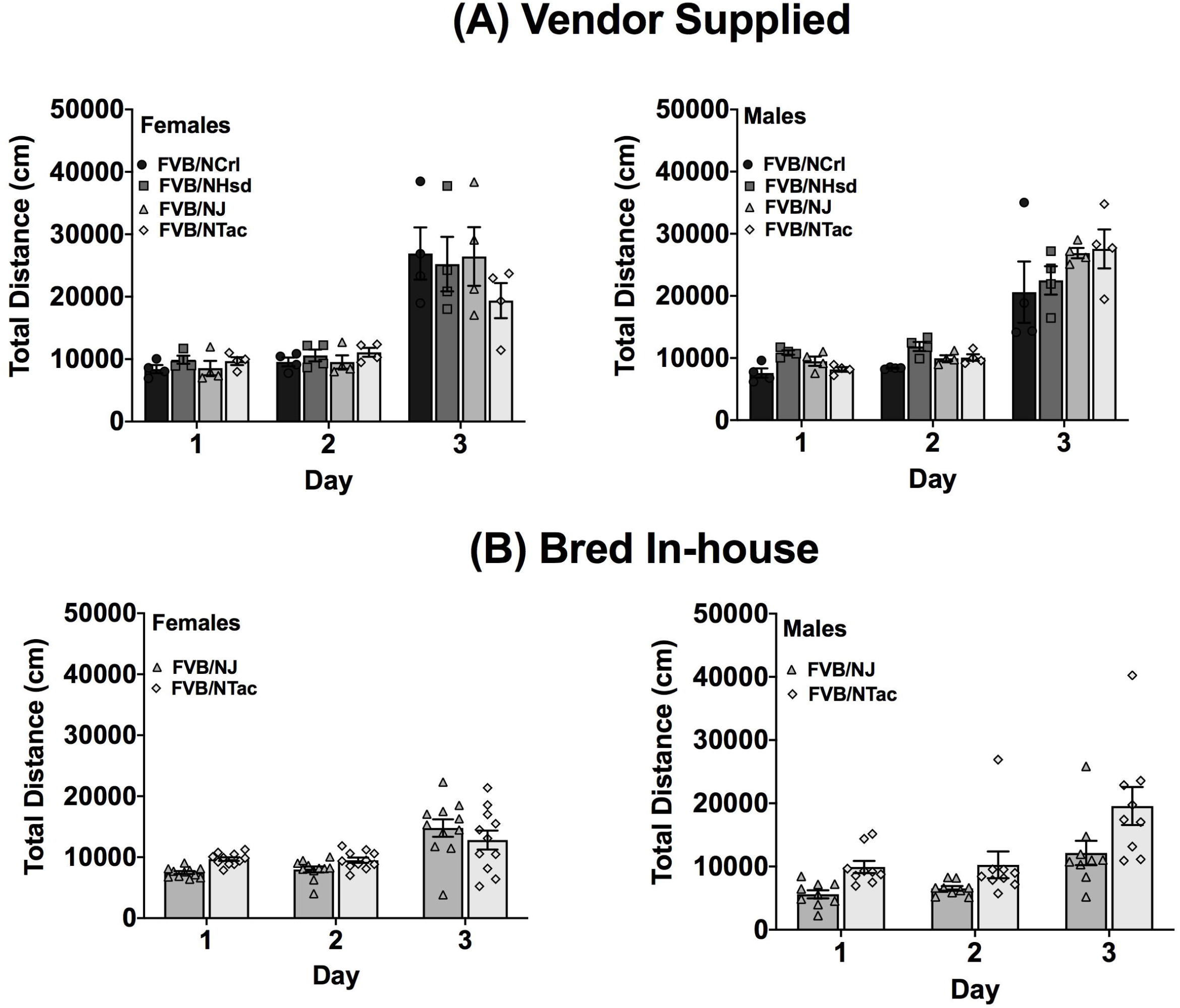
Acute cocaine induced locomotor activity for vendor supplied and bred in-house FVB/N substrains. (A) Total distance moved in 30-mins for each test day for vendor supplied female and male FVB/N substrains. All FVB/N substrains, regardless of sex, displayed significantly higher locomotor activity following cocaine administration on day 3. (B) Total distance moved in 30 mins for each test day for female and male FVB/N substrains bred in-house and cohoused at weaning. Overall, FVB/NTac male mice had significantly higher locomotor activity compared to FVB/NJ male mice. FVB/NJ female mice had higher cocaine induced locomotor activity compared to FVB/NTac female mice. Each data point represents an individual mouse and error bars are standard error of the mean.

#### Bred In-House

Locomotor activity in response to cocaine was significantly higher than response to saline in both of the FVB/N substrains that were bred in-house for 1-2 generations – FVB/NJ and FVB/NTac (F_(2,108)_ = 31.3; p = 1.9×10^−11^; **Fig 5B**). FVB/NTac mice exhibited higher locomotor activity across all three days in comparison to FVB/NJ mice (*t*(118) = −2.8; *p* = 0.007). There was no significant main effect of sex, but we did observe a significant strain x sex interaction. FVB/NTac males were significantly more active than FVB/NJ males (*t*(42) = −3.0; *p* = 0.005) but FVB/NTac and FVB/NJ females did not differ (*t*(64) = 0.60; *p* = 0.552).

### 3.6 Locomotor response to cocaine did not differ across 2 NOD substrains Bred In-House

NOD substrain testing was limited to those that were bred in-house at UNC. We observed a significant effect of day with both NOD/ShiLtJ and NOD/MrkTac showing a robust increase in locomotor activity on Day 3 after exposure to cocaine (**Fig 6**; F(_2,240_) = 177.9; *p* = 4.2 × 10^−48^). NOD/ShiLtJ mice were slightly less active on Days 1 and 2, but this difference was not significant (F_(1, 240)_ = 3.0; *p* = 0.084). No significant strain, sex or interaction effects were observed.

**Figure 6.**
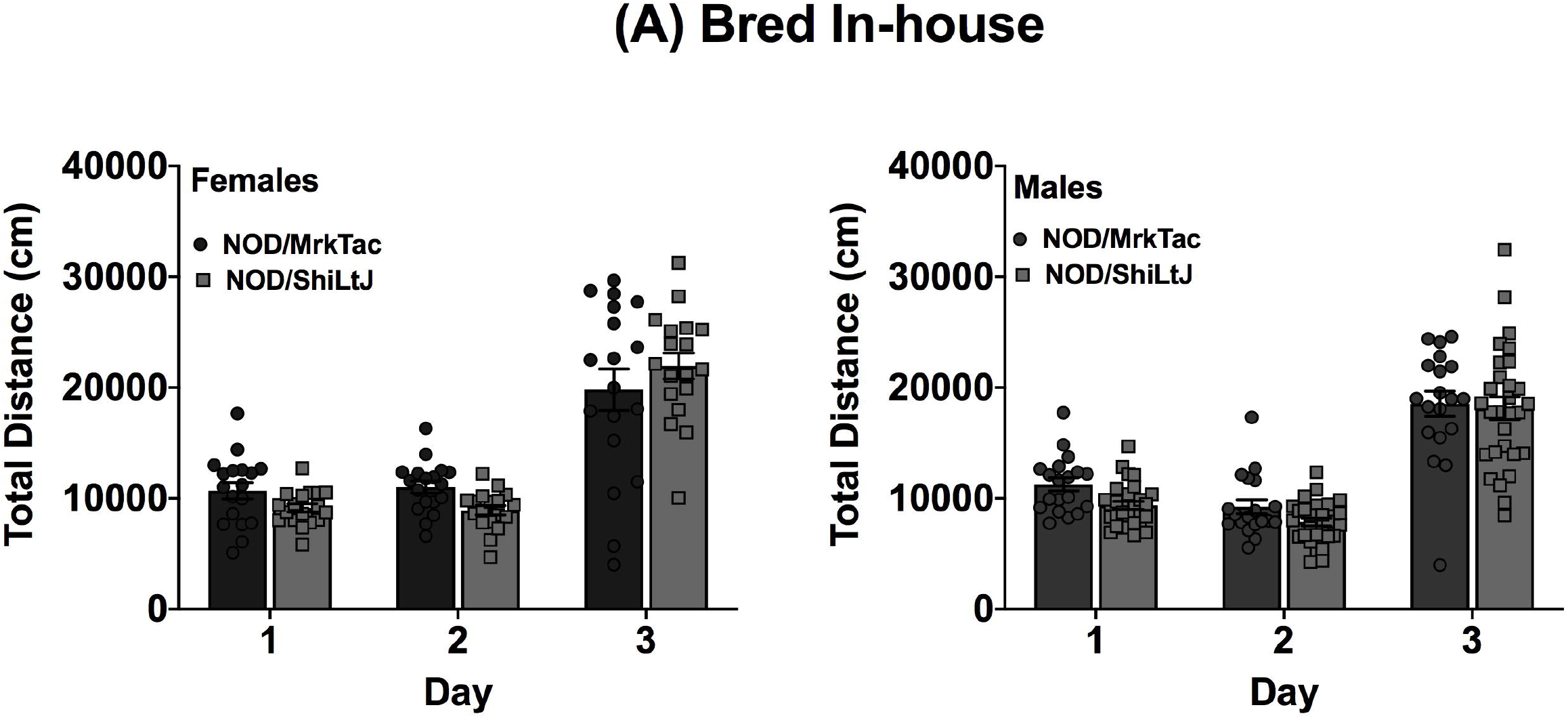
Acute cocaine induced locomotor activity for NOD substrains bred in-house. Total distance moved in 30 mins for each test day for female and male NOD substrains bred in-house. All mice, regardless of substrain, had higher locomotor activity following cocaine administration compared to saline. Each data point represents an individual mouse and error bars are standard error of the mean.

## 4.0 Discussion

Laboratory mice are invaluable tools in biomedical research and have contributed greatly to our understanding of biological and disease processes. Inbred strains, in particular, have been used for decades in genetic studies aimed at identifying genes and genetic loci that contribute to behavioral phenotypes including responses to various drugs of abuse (Tarantino, McClearn et al. 1998, Jones, Tarantino et al. 1999, Philip, Duvvuru et al. 2010, Yazdani, Parker et al. 2015). While these studies have been very successful in identifying chromosomal regions that likely harbor causal genetic variants, the genetic diversity present in mapping crosses between two standard inbred strains has hindered progress and these strategies have rarely progressed to identifying a specific causal gene or variant. The reduced genetic complexity in inbred mouse substrains offers the opportunity to overcome this hurdle and more rapidly and efficiently identify both the causative gene and specific genetic variant.

In order to use the RCC approach to identify causative genes and genetic variants one needs to identify substrains that exhibit phenotypic differences in the trait of interest. For example, this approach has been used successfully to identify the *Cyfip2* gene as a regulator of basal and cocaine-induced locomotor activity, behavioral sensitization and binge-eating in two C57BL/6 substrains, C57BL/6J and C57BL/6NJ (Kumar, Kim et al. 2013)(Kirkpatrick, Goldberg et al. 2017). The study of addiction-related behaviors, and specifically initial locomotor sensitivity to psychostimulants, has been mostly limited to the C57BL/6 inbred substrains. We assessed differences in cocaine-induced locomotor response across 20 inbred mouse substrains from 6 different strain sets. Thus, our data represent the first characterization of cocaine-induced locomotor activation in most of these substrains.

Two sets of strains showed particularly robust substrain differences that were replicated across experimental cohorts. C3H/HeNTac mice had significantly higher basal and cocaine-induced locomotor activity than C3H/HeNCrl and C3H/HeNHsd mice in one experimental cohort and C3H/HeJ mice in both cohorts (**Fig 3**). The similarity of the behavioral phenotype in C3H/HeJ, C3H/HeNCrl and C3H/HeNHsd substrains suggests that the causal variant or variants became fixed in the C3H/HeNTac substrain after it diverged from the other C3H/HeN lines in 1974 (C3H/HeNCrl) and 1983 (C3H/HeNHsd). However, we must also consider the possibility that C3H/HeNCrl and C3H/HeNHsd behavioral phenotypes result from a different variant or variants. We also observed consistent substrain differences in behavior in DBA/2 mice that were supplied by commercial vendors and and bred in-house. DBA/2NTac mice had significantly lower basal and cocaine-induced locomotor activity in comparison with DBA/2J and DBA/2NCrl in both cohorts (**Fig 4**). These data suggest that the causal variant(s) likely arose in the DBA/2NTac substrain after it diverged from DBA/2N in 1981.

Correspondence of significant substrain differences across the two cohorts suggests a strong genetic component and supports the use of the RCC to identify specific causal variants that influence basal locomotor activity and response to cocaine. We do note that we observed significant substrain differences in either basal or cocaine-induced locomotor behavior in 4 out of the 6 strain sets, suggesting that multiple genes likely contribute to these behaviors and more than one variant may be responsible for behavioral differences observed in any set of substrains. Therefore, larger mapping cohorts may be required for QTL identification (Bryant CD 2018), but the advantages of using a RCC for variant identification still apply.

We also identified different behavioral phenotypes in vendor-supplied substrains vs those bred in-house. For example, FVB/NTac mice bred in-house were significantly more active than FVB/NJ mice bred in-house across all three days of testing whereas vendor-supplied FVB substrains showed similar locomotor behavior across all three test days (**Figs 5A, B**). Neither A/J nor A/JOlaHsd substrains bred in-house were significantly activated in response to cocaine and in fact, locomotor behavior in these two substrains decreased significantly across all three days of testing (**Fig 1B**). Vendor-supplied A/J and A/JOlaHsd substrains were also not significantly activated in response to cocaine, but locomotor behavior was similar across test days with no significant decrease in activity (**Fig 1A**).

The observation of behavioral differences in the same substrain based on the source from which mice were obtained suggests that environmental factors could be responsible. Multiple studies have systematically examined environmental factors that might affect behavioral phenotypes including, but not limited to diet, type of cage, cage density, season, time of day, transportation and experimenter effects (Crabbe, Wahlsten et al. 1999, Chesler, Wilson et al. 2002, Chesler, Wilson et al. 2002, Wahlsten, Metten et al. 2003, Sorge, Martin et al. 2014). However, previous studies have generally assessed behavioral differences in mice tested across multiple sites. We examined behavior in all mice, independent of the source, in the same behavioral facility (and same testing room) at UNC. As such, we were able to control, to the extent possible, the environment to which the mice were exposed in the 5-week period leading up to testing. Mice were maintained on the same light cycle, tested during the same time of day, provided the same diet and water and housed in the same caging and animal holding room prior to and throughout testing.

The stress of transportation is an obvious difference between vendor-supplied mice and those bred in-house. We don’t believe transportation stress could fully explain observed behavioral differences between mice from different sources. Previous studies have shown that transportation has very little effect on behavioral outcomes (Crabbe, Wahlsten et al. 1999, Chesler, Wilson et al. 2002). Moreover, vendor-supplied mice arrived at UNC very close to weaning age and were habituated to our vivarium conditions for approximately 5 weeks prior to testing.

Experimenter effects may have also had an impact on behavioral outcomes. All mice supplied directly from the vendor were tested by the same animal handler (a female), whereas substrains bred in-house were tested by a group of 5 animal handlers including males and females. At least two studies have established that experimenter effects (Chesler, Wilson et al. 2002) and even the sex of the individual testing the mice (Sorge, Martin et al. 2014) can significantly affect the outcome of behavioral tests.

The gut microbiome has been implicated in numerous behavioral traits including locomotor response to psychostimulants (Kiraly, Walker et al. 2016, Meckel and Kiraly 2019). Composition of the gut microbiota, even in the same inbred strain background, can vary across time, from vendor to vendor, and even between different facilities and animal holding rooms at the same vendor or institution (Servick 2016, Ericsson and Franklin 2021). These differences can be attributed to a host of environmental factors including diet, caging, bedding and water supply (Lundberg, Bahl et al. 2017, Bidot, Ericsson et al. 2018, Ericsson, Gagliardi et al. 2018). Host genetic background also plays a significant role in the composition of the gut microbiota (Bubier, Chesler et al. 2021). Profound or even subtle changes in the gut microbiota in response to relocation from vendors to our vivarium could interact with different genetic backgrounds to significantly impact behavior. The relationship between genetic background and behavior becomes even more complicated when one considers that substrain behaviors attributable to stable differences in the gut microbiota could be erroneously ascribed solely to genetics. Shifts in the gut microbiota in response to changing environments could alter phenotypes and impact replicability from study to study. Recent studies have also established that the maternal microbiome can affect offspring neurodevelopment and impact behavior in adulthood (Codagnone, Stanton et al. 2019, Warner 2019, Vuong, Pronovost et al. 2020). Thus, it is important to consider not only the source of the mice being tested, but the composition of the maternal microbiome during neurodevelopment.

In summary, this study expands the knowledge of phenotypic differences in locomotor activity and initial response to cocaine in 6 sets of inbred mouse substrains which had previously not been characterized. Four of the six strain lineages displayed significant substrain differences in either basal- or cocaine-induced locomotor behavior and can be utilized in RCCs to identify causal genetic variants. Environmental factors also warrant follow-up, as differences in behavior were observed across the same inbred substrains obtained from different sources. C3H/He and DBA/2 lineages demonstrated stable and robust differences in cocaine-induced locomotor behavior, and are good candidates for additional studies to investigate genetic and environmental factors that contribute to initial cocaine sensitivity. Future studies can utilize these data to increase our understanding of the complex factors that increase CUD and potentially lead to new therapeutic targets.

