## Supplemental Table 1 for "Cocaine-induced locomotor activation differs across six sets of inbred mouse substrains"

| | | | | | | | | | Females | | | | | | Males | | | | | |
|---|---|---|---|---|---|---|---|---|---|---|---|---|---|---|---|---|---|---|---|---|
| Strain | Substrain | Vendor | Strain Catalog Number | Origin | Cage Environment | Female | Male | Total | Day 1 Mean | Day 1 StdDev | Day 2 Mean | Day 2 StdDev | Day 3 Mean | Day 3 StdDev | Day 1 Mean | Day 1 StdDev | Day 2 Mean | Day 2 StdDev | Day 3 Mean | Day 3 StdDev |
| A/J | A/J | Jackson Laboratory | #000646 | Bred in-house | Cohoused | 20 | 17 | 37 | 1432.8 | 891.1 | 991.8 | 764.3 | 523.1 | 451.2 | 1817.5 | 942.8 | 1247.8 | 756.9 | 903.9 | 1251.5 |
| | A/JOlaHsd | Envigo | #049 | Bred in-house | Cohoused | 19 | 17 | 36 | 824.8 | 597.2 | 383.4 | 472.2 | 190.6 | 390.4 | 1087.0 | 772.7 | 727.7 | 774.2 | 441.1 | 1029.9 |
| | A/J | Jackson Laboratory | #000646 | Vendor Supplied | Not cohoused | 8 | 8 | 16 | 1986.1 | 1030.5 | 1561.4 | 788.0 | 381.8 | 554.8 | 1125.5 | 659.5 | 1365.6 | 744.4 | 1312.2 | 1813.7 |
| | A/JCr | Charles River Laboratories | #563 | Vendor Supplied | Not cohoused | 8 | 8 | 16 | 1677.3 | 796.0 | 1778.3 | 860.7 | 3710.4 | 4200.6 | 1957.7 | 1044.4 | 1338.9 | 739.9 | 2759.4 | 4899.7 |
| | A/JOlaHsd | Envigo | #049 | Vendor Supplied | Not cohoused | 8 | 8 | 16 | 754.6 | 452.5 | 780.7 | 599.8 | 1244.1 | 2005.9 | 1032.9 | 580.1 | 1121.6 | 662.6 | 1261.1 | 1687.8 |
| BALB/c | BALB/cByJ | Jackson Laboratory | #001026 | Bred in-house | Not cohoused | 4 | 2 | 6 | 7217.4 | 3571.3 | 5257.9 | 2922.1 | 4278.9 | 3984.0 | 4709.8 | 29.6 | 81.8 | 46.7 | 880.0 | 810.5 |
| | BALB/cJ | Jackson Laboratory | #000651 | Bred in-house | Not cohoused | 6 | 0 | 6 | 6611.2 | 5270.8 | 5056.4 | 6427.4 | 5597.2 | 6249.3 | | | | | | |
| | BALB/cAnNCrl | Charles River Laboratories | #028 | Vendor Supplied | Not cohoused | 7 | 8 | 15 | 3378.9 | 1917.6 | 1363.6 | 1236.4 | 1312.0 | 1713.1 | 3641.1 | 1786.0 | 911.8 | 803.5 | 2770.1 | 4465.3 |
| | BALB/cAnNHsd | Envigo | #047 | Vendor Supplied | Not cohoused | 8 | 8 | 16 | 3893.3 | 1787.8 | 1982.2 | 1790.5 | 1784.7 | 1901.5 | 3017.1 | 1613.5 | 1542.8 | 1227.8 | 1511.0 | 3293.0 |
| | BALB/cByJ | Jackson Laboratory | #001026 | Vendor Supplied | Not cohoused | 7 | 8 | 15 | 7515.4 | 2737.0 | 3970.4 | 3288.1 | 10821.6 | 7551.5 | 3975.8 | 2669.8 | 3218.4 | 2961.9 | 7690.1 | 8439.3 |
| | BALB/cJ | Jackson Laboratory | #000651 | Vendor Supplied | Not cohoused | 8 | 8 | 16 | 6389.7 | 2109.3 | 4089.8 | 2679.2 | 6701.4 | 6789.9 | 5612.9 | 1781.9 | 2149.3 | 1844.8 | 2723.9 | 3765.6 |
| C3H/He | C3H/HeJ | Jackson Laboratory | #000659 | Bred in-house | Cohoused | 11 | 8 | 19 | 3305.0 | 810.6 | 3750.0 | 1603.1 | 18080.8 | 5727.2 | 3528.4 | 1490.3 | 3826.3 | 1641.8 | 18911.4 | 5873.4 |
| | C3H/HeNTac | Taconic | C3H-F/C3H-M | Bred in-house | Cohoused | 12 | 8 | 20 | 5669.8 | 831.2 | 6113.1 | 1603.7 | 36004.9 | 8427.3 | 5471.8 | 1854.7 | 5825.8 | 1964.7 | 26960.6 | 12045.8 |
| | C3H/HeJ | Jackson Laboratory | #000659 | Bred in-house | Not cohoused | 18 | 14 | 32 | 3398.7 | 1057.5 | 3511.2 | 1472.4 | 14617.3 | 5142.1 | 3498.3 | 852.1 | 3333.9 | 903.4 | 15314.5 | 6038.8 |
| | C3H/HeNTac | Taconic | C3H-F/C3H-M | Bred in-house | Not cohoused | 21 | 16 | 37 | 6019.4 | 2047.1 | 7199.9 | 2008.3 | 27739.8 | 11743.4 | 6905.3 | 1561.1 | 6720.5 | 1939.6 | 27654.4 | 10271.4 |
| | C3H/HeNHsd | Envigo | #040 | Bred in-house | Not cohoused | 32 | 32 | 64 | 4112.8 | 1432.8 | 4850.3 | 1469.5 | 19374.4 | 7991.4 | 3704.0 | 1677.0 | 3999.2 | 1716.9 | 14856.2 | 8853.0 |
| | C3H/HeNCrl | Charles River Laboratories | #025 | Bred in-house | Not cohoused | 19 | 32 | 51 | 3380.1 | 1763.3 | 3714.1 | 2018.8 | 17761.2 | 11014.1 | 3738.6 | 1293.0 | 4479.2 | 1203.3 | 19576.2 | 7152.1 |
| DBA/2 | DBA/2J | Jackson Laboratory | #000671 | Bred in-house | Cohoused | 20 | 18 | 38 | 5925.7 | 1120.5 | 5204.3 | 1959.3 | 13877.4 | 10429.2 | 5447.7 | 2101.7 | 4738.7 | 1931.9 | 12517.9 | 7447.5 |
| | DBA/2NCrl | Charles River Laboratories | #026 | Bred in-house | Cohoused | 11 | 19 | 30 | 6339.9 | 2085.8 | 5358.3 | 2046.3 | 15088.0 | 9565.1 | 6016.1 | 1122.3 | 4511.9 | 1370.8 | 11203.1 | 7388.3 |
| | DBA/2NTac | Taconic | DBA2-F/DBA2-M | Bred in-house | Cohoused | 19 | 15 | 34 | 5569.3 | 2062.3 | 3354.5 | 1759.7 | 6537.0 | 5796.6 | 6507.0 | 1475.8 | 4418.3 | 1336.9 | 5633.1 | 3263.9 |
| | DBA/2J | Jackson Laboratory | #000671 | Vendor Supplied | Not cohoused | 8 | 8 | 16 | 3787.3 | 2000.2 | 4129.7 | 2020.1 | 8043.1 | 6835.0 | 3264.8 | 1192.3 | 4900.4 | 788.0 | 12137.8 | 5662.2 |
| | DBA/2NCrl | Charles River Laboratories | #026 | Vendor Supplied | Not cohoused | 8 | 8 | 16 | 4543.2 | 2391.9 | 4179.7 | 2357.3 | 12003.3 | 12577.2 | 3560.4 | 1490.8 | 3132.5 | 1829.8 | 11248.2 | 4662.4 |
| | DBA/2NTac | Taconic | DBA2-F/DBA2-M | Vendor Supplied | Not cohoused | 8 | 8 | 16 | 2519.2 | 1805.3 | 1426.6 | 1263.5 | 4456.6 | 4490.4 | 2145.4 | 2331.9 | 1580.9 | 1743.0 | 2178.1 | 2156.5 |
| FVB/N | FVB/NJ | Jackson Laboratory | #001800 | Bred in-house | Cohoused | 11 | 9 | 20 | 7476.6 | 797.6 | 7990.5 | 1636.5 | 14784.8 | 4757.5 | 5606.9 | 1921.9 | 6537.1 | 1120.4 | 12139.6 | 5758.2 |
| | FVB/NTac | Taconic | FVB-F/FVB-M | Bred in-house | Cohoused | 11 | 9 | 20 | 9687.1 | 972.8 | 9504.4 | 1436.5 | 12803.6 | 5148.7 | 9931.8 | 2883.1 | 10260.0 | 6354.3 | 19566.4 | 9029.5 |
| | FVB/NCrl | Charles River Laboratories | #207 | Vendor Supplied | Not cohoused | 4 | 4 | 8 | 8405.1 | 1304.0 | 9549.8 | 1398.9 | 26917.5 | 8366.8 | 7585.5 | 1498.5 | 8372.8 | 143.6 | 20593.8 | 9867.8 |
| | FVB/NHsd | Envigo | #118 | Vendor Supplied | Not cohoused | 4 | 4 | 8 | 9899.0 | 1290.6 | 10596.0 | 1884.7 | 25221.5 | 8730.3 | 10881.7 | 694.4 | 11859.5 | 1448.5 | 22500.1 | 4567.3 |
| | FVB/NJ | Jackson Laboratory | #001800 | Vendor Supplied | Not cohoused | 4 | 4 | 8 | 8562.1 | 2309.6 | 9537.4 | 2154.4 | 26447.7 | 9389.4 | 9488.7 | 1510.0 | 9968.9 | 929.0 | 26892.3 | 1669.2 |
| | FVB/NTac | Taconic | FVB-F/FVB-M | Vendor Supplied | Not cohoused | 4 | 4 | 8 | 9682.4 | 1257.9 | 11098.1 | 1430.5 | 19374.7 | 5623.9 | 8162.8 | 733.0 | 10074.8 | 1049.7 | 27565.5 | 6272.4 |
| NOD | NOD/MrkTac | Taconic | NOD-F/NOD-M | Bred in-house | Cohoused | 18 | 19 | 37 | 10695.1 | 3146.5 | 11021.8 | 2319.8 | 19816.5 | 7952.8 | 11257.3 | 2545.6 | 9242.7 | 2770.4 | 18533.8 | 4969.6 |
| | NOD/ShiLtJ | Jackson Laboratory | #001976 | Bred in-house | Cohoused | 18 | 29 | 47 | 9133.8 | 1561.9 | 8923.1 | 1766.1 | 21956.3 | 4947.3 | 9369.3 | 1937.1 | 7865.4 | 1886.6 | 18134.8 | 5490.8 |
